# Critical Process Steps for Mechanical Agitation Driven Coated Nanobubble Self Assembly

**DOI:** 10.64898/2026.07.20.736752

**Authors:** Theresa Kosmides, Dana Wegierak, Aaqib H. Khan, Ilya Bederman, Tae Kyong John Kim, Agata A. Exner

## Abstract

The term nanobubble (NB) includes a wide range of gas core, submicron particles. A subgroup of NBs consists of phospholipid-shelled (or coated) nanoparticles stabilizing a perfluorocarbon gas core which have gained recent interest as ultrasound (US) contrast agents. Several methods are available to produce coated NBs. Among these, amalgamation driven self-assembly has been the most utilized. Amalgamation (also referred to as mechanical agitation) is a simple technique currently used for production of commercial and clinically relevant microbubble suspensions. When combined with size-isolation steps, it can also generate submicron NB suspensions with a narrow size distribution. While this technique has been used extensively, no prior work has systematically examined the critical manufacturing parameters needed to produce the optimal coated NB formulation. In this work, we investigate how the precursor lipid dispersion, perfluorocarbon gas to lipid ratio, and pressurized size isolation affect the formation and size isolation of stable, uniform NBs. Results show that the precursor lipid dispersions exhibiting a monomodal size distribution produced the most stable NBs. Additionally, perfluorocarbon volume in excess of lipid dispersion volume is required to form high concentration, stable NBs. Finally, pressurized size isolation resulted in high concentration, US stable NBs. These findings establish the understanding of the key process parameters which affect uniform size and stable NB production via mechanical amalgamation.

## Introduction

Nanobubbles (NBs) are nanoparticles with a gas core and diameter less than 1000 nm.^1^ This definition encapsulates a wide range of submicron particles including surface NBs (surface bound, gas-filled cavities), bulk NBs (spherical uncoated gas particles in suspension), and coated NBs (polymer, lipid, or protein shell stabilized).^1–3^ The name NB is often used to refer to ultrafine bubbles, which are ISO recognized as a “gas in a medium enclosed by an interface with a volume equivalent diameter of less than 1 µm”.^4^ These ranging definitions cover the extremely wide array of NBs and their unique applications in fields such as water treatment, agriculture, and medicine.^5 – 7^ As the field of NB research is expansive, in this work we will focus on submicron, phospholipid shell stabilized, perfluoropropane (C_3_F_8_) core NBs or specifically, coated, bulk, ultrafine bubbles, herein referred to as NBs.

Lipid-shelled NBs with a mean diameter of 200 – 400 nm, have been investigated in numerous biomedical applications, but a primary area of interest is their use as an ultrasound (US) contrast agent and US-triggered drug delivery system.^8–10^ While clinically utilized microbubble (MB) agents are used routinely as blood-pool enhancing agents in echocardiography, liver, kidney and breast lesion detection^11– 13^, the NB submicron diameter allows them to move beyond the vascular endothelium in states of hyperpermeability, making them ideal ultrasound contrast agents (UCAs) for diagnostic imaging in extravascula r applications such as oncology and type 1 diabetes.^8,14,15^

Currently, NBs are used as exclusively preclinical UCAs; however, there has been recent progress in large animal studies towards eventual clinical translation.^16,17^ As the field advances, it is paramount to understand the critical manufacturing parameters for NBs to support translation. A thorough understanding of how NBs are affected by each manufacturing step will allow for future scaling to production scale and repeatable, reproducible agent production; a critical aspect of successful clinical translation. Generally, the focus of NB formulation research has been on developing novel formulations (i.e. ligand, fluorophore, antibody addition) rather than on evaluating the impact of each processing step on the final NB product.^9,18,19^ For the successful use of NBs as clinically relevant UCAs, they must be submicron in diameter, highly concentrated (>10^10^ NB/mL), and stable both in and out of the presence of US as US is inherently destructive to UCAs. To achieve this, the components of UCAs are carefully selected. Perfluorocarbons (PFCs) are commonly used in the core as they are relatively insoluble in water, increasing NB stability through slower gas diffusion. Additionally, shell components can be altered to increase stability; additives can be introduced to modify membrane stiffness, and polymer or protein shells can increase shell density compared to phospholipid shells.

Fundamentally, lipid-shelled NBs are formed by stimulus driven self-assembly and bubble nucleation. There are many methods to form NBs via self-assembly, including sonication, microfluidics, extrusion and mechanical amalgamation.^20 – 2 3^ Mechanical amalgamation driven self-assembly is a technically straight forward, low cost technique that is currently used to produce on market MBs.^24^ It is a process in which a lipid dispersion in an aqueous solvent serves as a precursor which, when coupled with a PFC gas headspace, is agitated at a high frequency (~76 Hz) typically using a repurposed dental amalgamator to initiate bubble self-assembly.^23,25,26^ The phospholipids in the lipid dispersion self-assemble into a monolayer around nucleated gas inclusions causing their hydrophobic chains to orient towards the bubble core and the hydrophilic polar head groups to the outer bubble shell.^27^ This process creates a highly polydisperse bubble population (diameter: ~100 – 5000 nm) from which NBs are isolated via differential centrifugation which results in low polydispersity (mean diameter: ~200 – 400 nm) and high concentration. While the applications of NBs produced via mechanical amalgamation have been well documented, a robust investigation into the critical processing steps to produce highly concentrated, stable NBs with uniform size has not been conducted.

To produce NBs via mechanical amalgamation, an oil-in-water lipid dispersion is first created by sonicating lipids solubilized in propylene glycol into a PBS/glycerol solution (Fig. 1A). Next, the suspension is degassed, the perfluorocarbon gas which makes up the core is added, and bubbles are formed via mechanical amalgamation driven self-assembly (Fig. 1B), then NBs are isolated via differential centrifugation (Fig. 1C). Finally, to evaluate if the NBs produced are echogenic (e.g. producing detectable acoustic activity), NBs are imaged in a tissue mimicking phantom using a preclinical ultrasound system (Fig. S1). In this work, we present a systematic analysis of critical NB manufacturing steps via in-house NB solutions. Specifically, we investigate three critical NB manufacturing steps: (1) size distribution of particles in the lipid dispersion precursor (shown throughout in blue), (2) ratio of perfluorocarbon gas to lipid dispersion (shown in green), and (3) pressurized size isolation (shown in red), and evaluate their effects on NB physical characteristics, stability and ultrasound responsiveness.

**Fig. 1.**
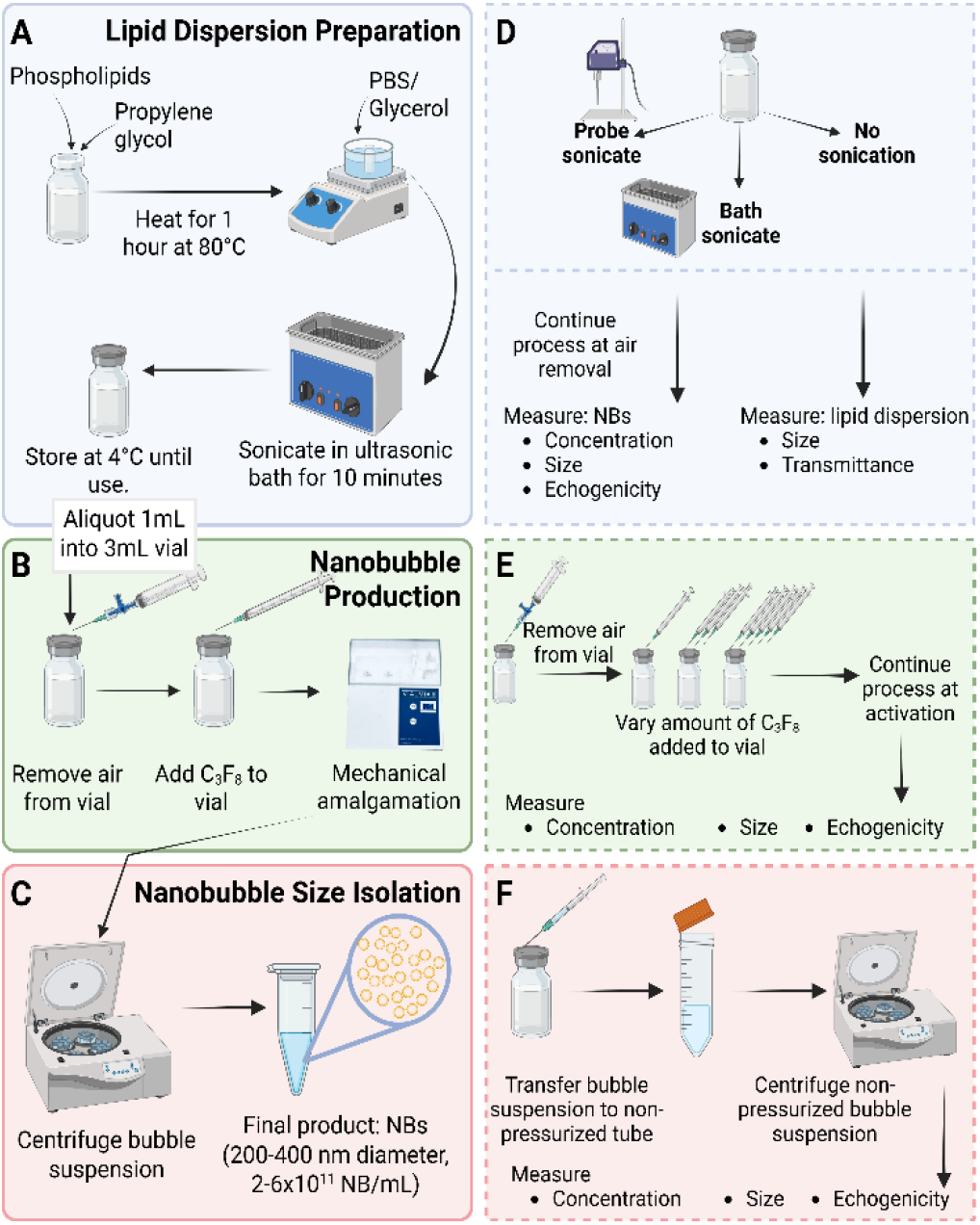
Overview of lipid dispersion, nanobubble preparation and size isolation process showing lipids are dissolved in propylene glycol (oil phase) prior to PBS, glycerol (water phase) addition to produce the lipid dispersion (a), the headspace in a sealed vial is replaced with perfluorocarbon and a polydisperse bubble suspension is created by amalgamation (b), nanobubbles are isolated via differential centrifugation resulting in a highly concentrated, submicron population (c), investigation of sonication intensity’s role on lipid dispersion and resulting NBs (d), investigation of gas to lipid ration on resulting NBs (e), investigation of pressurized centrifugation on resulting NBs (f).

## Experimental

### Preparation and isolation of nanobubbles

NBs were prepared as previously reported.^23^ Briefly, phospholipids (1,2-dibehenoyl-s n-gl ycero-3-phosphocholine (DBPC), 1,2-dipalmitoyl-s n-gl ycero-3-phosphate (DPPA), 1,2-dipalmitoyl-sn-glycero-3-phosphoethanolamine (DPPE)) (Avanti Polar Lipids, Alabaster, AL, USA) and 1,2-distearoyl-snglycer o-3-phosphoethanolamine-N-[methoxy(polyethyleneglycol)-2000] (ammonium salt) (mPEG-DSPE) (Laysan Bio, Arab, AL, USA) were dissolved in propylene glycol (PG) (Sigma Aldrich, St. Louis, MO, USA) at 80°C. Following dissolution, phosphate buffered saline (PBS) (Gibco Inc., Billings, MT, USA), and glycerol (Thermo Scientific Chemicals, Waltham, MA, USA) were added to create the phospholipid dispersion. The dispersion was sonicated in a bath sonicator for 10 minutes at room temperature. 1 mL of the dispersion was transferred to a 3 mL vial and headspace was replaced by C_3_F_8_ and mechanically amalgamated via VialMix for 45 seconds (Bristol-Myers Squibb Medical Imaging Inc., N. Billerica, MA, USA). NBs were isolated by differential centrifugation at 50 g for 5 minutes in an inverted vial. After centrifugation, 400 µL of NB solution was removed, comprising the stock sample.

To test the effect of sonication intensity, following the addition of aqueous (PBS, glycerol) to oil (lipids/PG) phase, the lipid dispersion underwent bath sonication (control) for 10 minutes, probe sonication (QSonica Q125, 30% amplitude, 30 s on/2 s off), or no sonication (Fig. 1D). The bath sonication was conducted at room temperature. The sample which underwent probe sonication was kept on ice during the treatment to avoid sample heating caused by probe sonication. The initial protocol was then followed as previously described beginning at aliquoting step.

To assess the role of atmospheric air contamination in the vial headspace, initial protocol was followed until headspace replacement. Instead, the amount of PFC (C_3_F_8_) added to the vial was controlled, resulting in gas to lipid ratios of 1:1, 3:1, 5:1 (v:v) (Fig. 1E). To achieve this, no headspace was removed after sealing the vial. When the C_3_F_8_ was added to the vial, the vial was not vented to control the volume of gas added to the system. Following C_3_F_8_ addition, the process followed the protocol as initially described.

For experiments to determine the role of a pressurized vessel on NB size isolation, the initial protocol (bath sonication, C_3_F_8_ in excess of lipid dispersion) was followed until differential centrifugation. For non-pressurized size isolation, after mechanical agitation, the material was transferred from the pressurized, gas tight vial to a falcon tube with atmospheric air in the headspace (Fig. 1F). NBs were size isolated via differential centrifugation at 50 g for 5 minutes. After centrifugation, 400 µL of NBs were removed.

### Size and transmittance of lipid dispersion

Precursor phospholipid dispersion particle size was determined by dynamic light scattering (DLS) with a 658 nm light source (LiteSizer 500, Anton Paar, Graz, Austria). The size distributions are reported from intensity-weighted (favors large particles) and number-weighted (favors small particles) analyses. The transmittance of light through each lipid dispersion was also determined by DLS. All DLS samples were diluted 1:1000 in PBS and measured at 25°C in triplicate.

### Size and concentration of nanobubbles

NB size was determined via DLS and resonant mass measurement (RMM) (Archimedes, Malvern Panalytical Inc., Westborough, MA, USA). NB concentration was determined by RMM. RMM is a technique which determines mean nanoparticle size, size distribution, and concentration by measuring the frequency changes in a resonating cantilever. Positively buoyant nanoparticles (NBs) will increase the frequency while negatively buoyant nanoparticles will decrease the frequency of the cantilever. The magnitude of the frequency shift relative to calibration beads is used to determine the size of the nanoparticles.

### X-ray photoelectron spectroscopy (XPS)

Elemental analysis of lipid dispersion and NBs was performed with a PHI Versaprobe 5000 Scanning X-ray Photoelec tron Spectrometer (XPS). Samples (10 µL) were freeze-dried on silicon wafers and blown dry with argon gas. XPS spectra were obtained using a monochromatic Al Kα X-ray beam. A sample spot size of 200 x 200 µm was used. Chemical composition was determined with built-in software (Multipak) and binding energies were referenced relative to the C1s peak at 284.5 eV. All samples were measured in triplicate.

### Gas chromatography and mass spectrometry

To confirm the presence of encapsulated C_3_F_8_ gas, 500 µL of NB were sealed in gas chromatography-mass spectrometry (GC-MS) vials. To facilitate gas release into the headspace, samples were sonicated in a 50 °C water bath for 20 minutes prior to analysis. Gas quantification was performed using GC-MS (Agilent Technologies, Santa Clara, CA, USA) consisting of a 5977B mass spectrometer coupled to a 7890B gas chromatograph. Briefly, 1 μL of sample was injected at a 1:20 split ratio with an injector temperature of 200 °C. Separation was achieved using an HP-5MS capillary column (30 m × 250 μm × 0.25 μm) maintained under a helium flow rate of 1.5 mL/min. The oven temperature was initially held at 60 °C for 1 min, followed by a ramp of 30 °C/min to 120 °C and held for an additional 5 min. The transfer line and detector temperatures were maintained at 250 °C. Analyses were performed in selected ion monitoring (SIM) mode with an electron multiplier gain factor of 2, where characteristic C_3_F_8_ fragments at m/z 69 (CF_3_+) and m/z 169 (C_2_F_5_+) were monitored. The dominant CF_3_ + fragment was quantified at a retention time of 4.48 min, and all detected fragments were confirmed using the NIST mass spectral database. A calibration curve was generated using known quantities of C_3_F_8_ gas to correlate peak area with gas volume, thereby enabling quantification of the equivalent gas volume present in the headspace of unknown samples.

### Echogenicity of nanobubbles

B-mode and nonlinear contrast US imaging was conducted with a Vevo 2100 preclinical US imaging system and MS250 transducer (FUJIFILM VisualSonics, Toronto, Canada) with a 21 MHz center frequency (f_c_) and bandwidth of 13-24 MHz. Images were acquired with the following parameters: 18 MHz transmit frequency, 4% power, 1 fps, 35 dB contrast gain and 18 dB 2D gain. NBs were suspended in PBS at 1×10^8^ NB/mL (based on initial NB concentration as determined by RMM) to capture initial signal intensity and signal decay. Diluted NBs were placed in a thin channel agarose phantom (1.5% agarose in deionized water) for imaging. The channel dimensions are equivalent to the transducer element dimensions (22 × 1 × 10 mm) and the transducer was directly coupled to the sample, ensuring the entire NB suspension was exposed to US during image acquisition. Two types of acquisitions were performed to investigate the stability of NBs in the presence of US (continuous) or without the presence of US (interval). For continuous imaging, 310 frames (approximately five minutes) were collected. For interval imaging, three frames were collected every 30 seconds for five minutes. For all acquisitions, the nonlinear contrast signal intensity at the focal depth was calculated via MATLAB (Mathworks, Natick, MA, USA) post processing.

## Results and Discussion

### Sonication effect on lipid dispersion precursor

To investigate the role of the size distribution of the lipid dispersion precursor on the resulting NBs, the dispersion was exposed to varying levels of sonication: no sonication, bath sonication (~0.03-0.1 W/cm^2^), or probe sonication (~100-700 W/cm^2^), and the resulting particles were analyzed with DLS, RMM and XPS. The dispersion size decreased with increasing sonication intensity (none, bath, probe, respectively) resulting in a 30x decrease in hydrodynamic diameter (HD) from no sonication to bath sonication and 36x decrease in HD from no sonication to probe sonication (Fig. 2A, B). The precursor particles became increasingly monomodal with increasing sonication intensity as seen in the 1.6x (bath sonication) and 2.1x (probe sonication) reduction in the polydispersity index compared to no sonication (Fig. 2A, B). With increasing sonication intensity, the optical transmittance of the dispersion significantly increased (mean ± SD: 52E-3 ± 35E-5, 59E-3 ± 21E-3, 13 ± 29E-2) (Fig. 2C). Finally, the elemental composition of the lipid dispersion precursor as determined by XPS did not significantly vary across sonication conditions (Fig. 2D). This data suggests that increasing sonication intensity results in more uniform size and smaller precursor particles in the lipid dispersion precursor without altering the elemental composition of the dispersion.

**Fig. 2.**
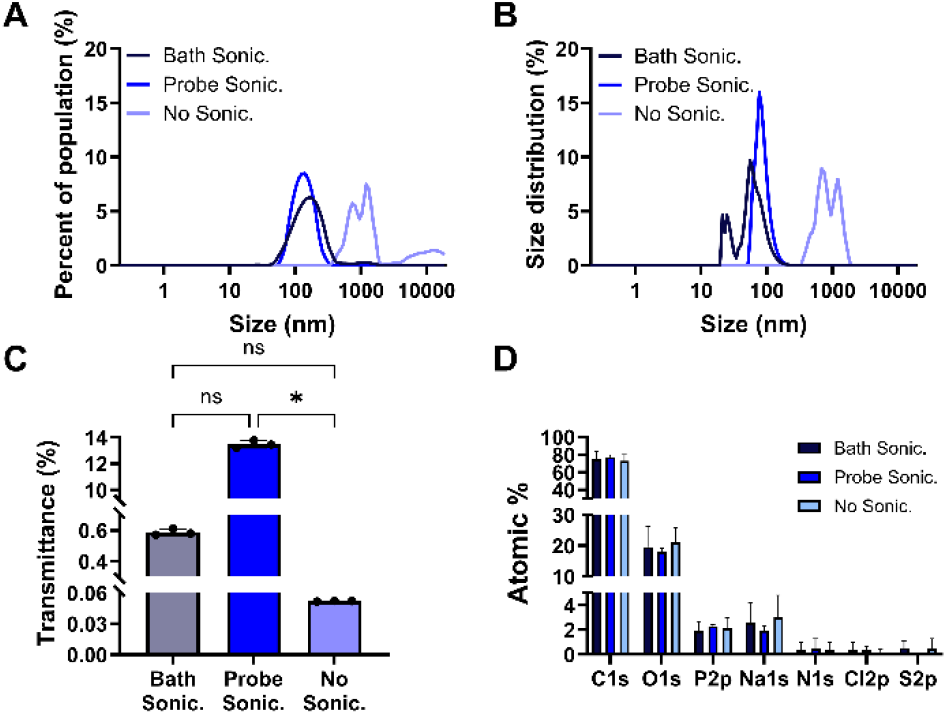
Characterization of lipid dispersion precursor material which underwent variable sonication conditions showing intensity weighted mean DLS size distribution of lipid dispersions (a), number weighted mean DLS size distribution of lipid dispersions (b), light transmittance through lipid dispersion precursor dispersions (c), XPS elemental composition of lipid dispersions (d). All samples are n=3.

Surprisingly, NBs generated from no sonication, bath sonicated, or probe sonicated lipid dispersions, had no significant difference in size (mean ± SD: 231.0 ± 6.9, 230.9 ± 5.0, 239.5 ± 12.6 nm) or concentration (mean ± SD: 46.2E11 ± 96.9E 9, 44.7E11 ± 44.1E9, 44.3E11 ± 53.3E9 NB/mL) regardless of lipid precursor conditions, respectively (Fig. 3A-D). NBs produced with probe sonicated lipid dispersion had the lowest fraction of non-buoyant particles as measured through RMM (Fig. 3D); though the difference was not statistically significant. Likewise, there was no significant difference in the initial nonlinear signal intensity for all samples, however the probe sonicated NBs were more stable and showed a slower decay in acoustic activity decayed compared to the bath and no sonication conditions (Fig. 3E, F). As previously noted, the probe sonicated NBs had the smallest fraction of non-buoyant particles (e.g. micelles, liposomes). The non-buoyant particles do not contribute to US signal generation and therefore the higher fraction of NBs is likely the cause of the increased stability in the presence of US.

**Fig. 3.**
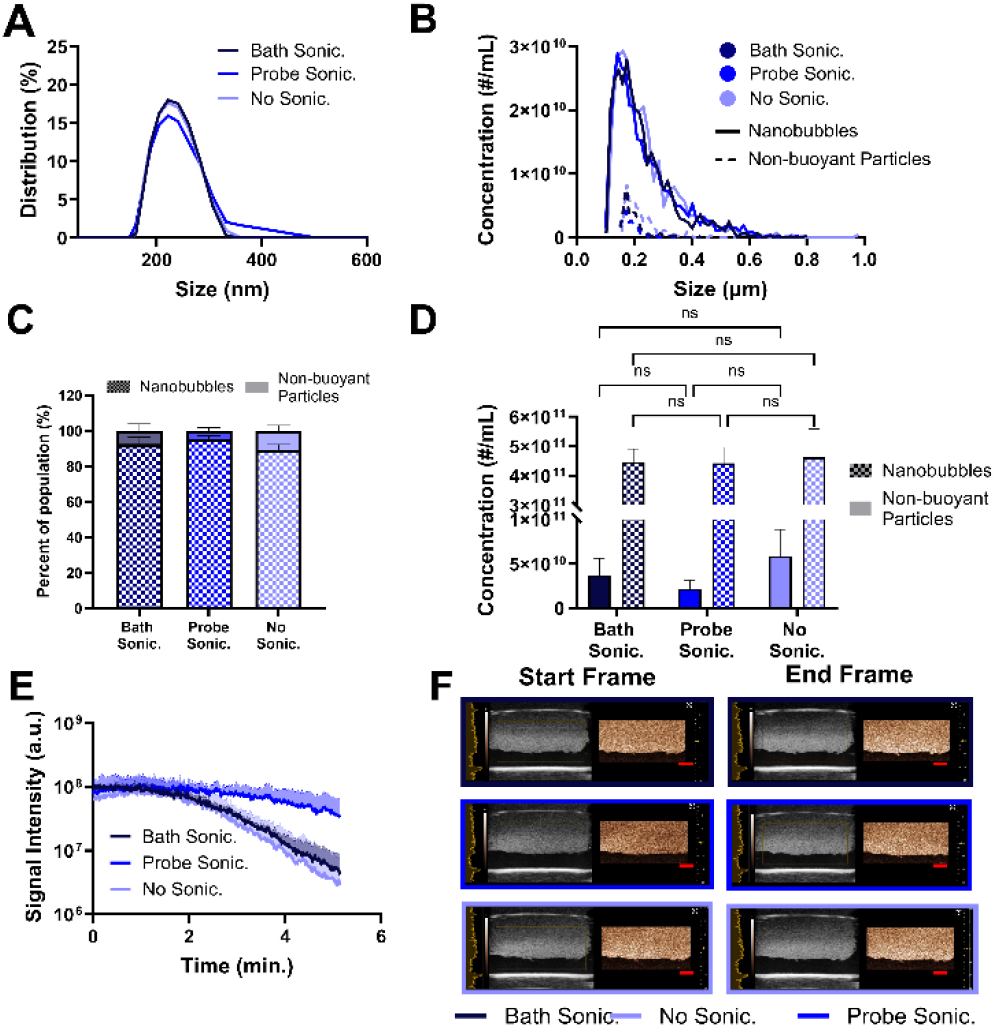
Characterization of nanobubbles made from lipid dispersions which underwent variable sonication conditions showing mean intensity DLS size distributions of NBs (a), mean RMM concentration distributions of NBs and non-buoyant particles (b), percent of sample which are nanobubbles vs non-buoyant particles (c), mean nanobubble vs non-buoyant particle concentration (d), mean NLC US signal intensity of concentration matched nanobubble samples (e), representative start and end frame B-mode (left, grayscale) and NLC mode (right, copper) US images of concentration matched nanobubble samples (f). All samples are n=3. Scale bar is 3 mm.

### Effect of perfluorocarbon gas to lipid volume on nanobubbles

NBs were produced while increasing the ratio of C_3_F_8_ gas to lipid dispersion precursor from 1:1, to 3:1 and 5:1 (v:v). The resulting NBs had similar size distributions across both number and intensity weighted averages for all conditions (mean ± SD: intensity 269.4 ± 13.2, 244.9 ± 3.5, 239.6 ± 3.6) (Fig. 4A). The number weighted average is more sensitive to smaller particle fractions while the intensity weighted average tends towards large particles; thus, when there is agreement between these values, it suggests that the sample has a uniform size distribution and there are no detectable aggregates or large particles. When the C_3_F_8_ gas was equal in volume to the lipid dispersion, the resulting NB concentration decreased by an order of magnitude as compared to both conditions of NBs produced with excess PFC (Fig. 4B). Despite the lower concentration, the resulting size distribution of the NBs was similar to that of NBs produced with higher C_3_F_8_ gas volumes. The NBs produced with a 1:1 ratio had the largest non-buoyant particle fraction compared to the excess C_3_F_8_ conditions; 4.3x higher than 3:1 and 3.5x higher than 5:1 (Fig. 4C). NBs produced with a 3:1 ratio of C_3_F_8_ gas to lipid dispersion had a significantly higher concentration (1.4x) of NBs but no significant difference in non-buoyant particles compared to the NBs produced with a 5:1 ratio (Fig. 4D).

**Fig. 4.**
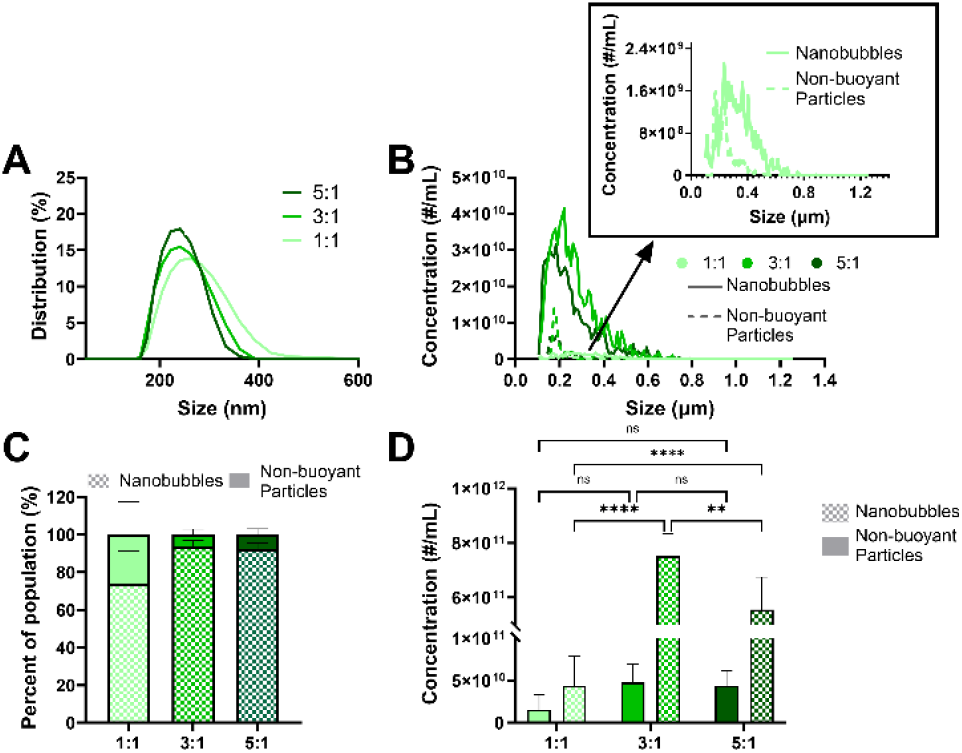
Characterization of nanobubbles produced with varying volumes of C_3_F_8_ gas showing mean intensity weighted DLS size distribution of nanobubbles made from varying ratios of C_3_F_8_ to lipid dispersion (a), mean RMM concentration distributions of nanobubbles made from varying ratios of C_3_F_8_ to lipid dispersion (b), percent of sample which is nanobubble vs non-buoyant particle (c), mean nanobubble vs non-buoyant particle concentration (d). All samples are n=3.

Overall, the data show that excess C_3_F_8_ gas volume is required to produce high concentration (>10^11^ #/mL) NBs. Excess PFC volume may increase the proportion of dissolved PFC in the solution, thus reducing the driving force for outward gas diffusion from the NB core. This effectively improves NB stability and US signal intensity by reducing outward gas diffusion, a critical component of NB US signal generation. The ratio is not critical to the size distribution of the resulting NBs; however it is critical to NB concentration. These findings suggest that when there is insufficient gas volume during the amalgamation driven self-assembly, the proportion of gas core (i.e. NBs) decreases while the proportion of non-gas core particles (i.e. micelles, liposomes), which do not produce US signal, increases.

### Effect of perfluorocarbon volume on nanobubble stability under ultrasound

Under continuous US, the 5:1 NB signal intensity decreased by approximately 10x while the 3:1 and 1:1 NBs decayed almost 100x across 5 minutes (Fig. 5A). Without the presence of continuous US, the 5:1 and 3:1 NB signal intensity remained approximately stable, with no significant signal loss while the 1:1 NBs had a 4x signal decrease (Fig. 5B). The 5:1 NBs had significantly higher initial signal compared to the 3:1 (4.5x) and 1:1 (13x) NBs (Fig. 5C). The decay in NB acoustic activity can be observed visually in the loss of signal (copper/white speckle) from the start to end frame (Fig. 5D). The higher initial NLC signal intensity for the 3:1 and 5:1 NBs may be due to a higher proportion of C_3_F_8_ in the NB core. Additionally, the significantly higher proportion of non-buoyant (unable to generate NLC US signal) particles in the 1:1 NBs likely contribute to lower initial signal intensity. Both the 3:1 and 5:1 NB conditions had significantly higher PFC than the 1:1 NB condition, 58x and 75x respectively (Fig. 5E). The 5:1 NB condition has the highest initial NLC signal but no significant difference in concentration; suggesting that the increased signal intensity results from a larger proportion of C_3_F_8_ in the NB cores.

**Fig. 5.**
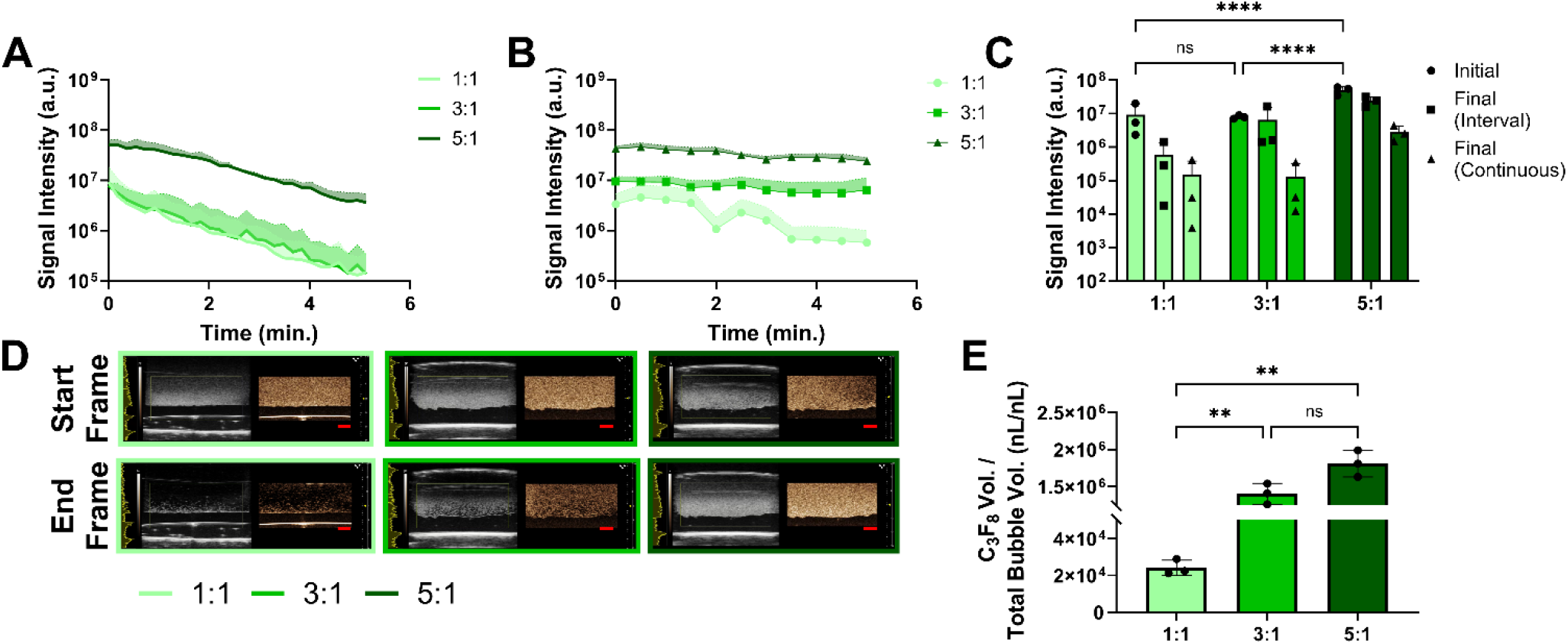
Echogenicity characterization of nanobubbles produced with varying volumes of C_3_F_8_ gas showing continuous acquisition ultrasound data (a), interval acquisition ultrasound data (b), comparison of initial and final signal intensity from continuous and interval acquisitions (c), representative B mode and NLC mode start and end frames of the continuous acquisition samples (d), gas headspace analysis of NBs of C_3_F_8_ to bubble volume ratios (e). All samples are n=3. Scale bar = 3 mm.

In contrast to the continuous acquisition, the interval acquisition subjects the NBs to ~89% less US energy; energy that can inertially cavitate, or destroy, the NBs. Despite this significant decrease in applied energy, the NBs with equal PFC to lipid volume demonstrated a significant decrease in US signal; suggesting inherent instability. This is likely due to atmospheric air contamination in the NB core. Atmospheric gases (i.e. N_2_, O_2_, Ar, CO_2_) have a high diffusivity, resulting in diffusion outward from the NB core and thus reduced US signal generation and bubble stability.

These findings suggest that the stability of NBs both with and without the presence of US can be tuned via gas to lipid ratio adjustment. These parameters can be translated directly to increased experimental control and improved clinical signal acquisition. In some cases, the time lost to agent instability is the critical consideration of a procedure’s feasibility (i.e. pharmacokinetic studies).

### Effect of pressurized size isolation on nanobubbles

Following mechanical amalgamation, NBs were isolated by centrifugation either in the original sealed and degassed vial or in a falcon tube at atmospheric pressure. There was no significant difference in size of NBs size isolated with either method (Fig. 6A, B). The agreement in size across both intensity- and number-weighted averages suggests that the resulting populations are uniform. There was also no significant difference in the proportion of NBs to non-buoyant particles between pressurized and non-pressurized (Fig. 6C). However, the NBs that were size isolated via non-pressurized size isolation had significantly lower concentration (0.68x) compared to pressurized size isolation (Fig. 6D). In the presence of US, the pressurized NBs resulted in high nonlinear signal intensity throughout the 5 minute acquisition and less variable signal decay compared to non-pressurized NBs (Fig. 6E). Both samples had minimal NLC signal intensity decay, as demonstrated by the visually negligible change in copper/white speckle in the NLC image (Fig. 6F).

**Fig. 6.**
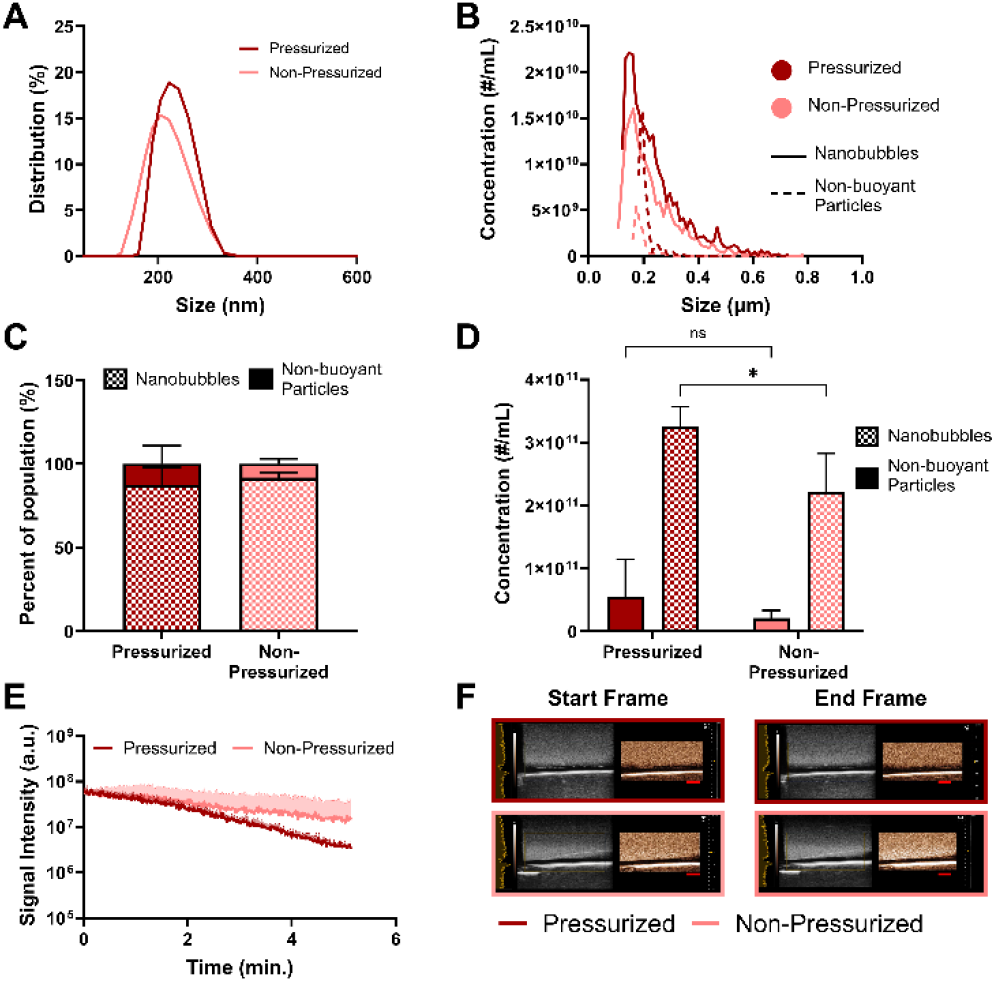
Characterization of nanobubbles size isolated via pressurized or non-pressurized centrifugation showing mean intensity DLS size distribution of nanobubbles (a), mean RMM concentration distributions of nanobubbles and non-buoyant particles (b), percent of sample which is nanobubble vs non-buoyant particles (c), mean nanobubble vs non-buoyant particle concentration (d), nonlinear contrast ultrasound signal intensity of concentration matched nanobubble samples (e), representative start and end frame B-mode and NLC mode US images of concentration matched nanobubble samples (f). All samples are n = 3. Scale bar is 3 mm.

Pressurization during centrifugation does not significantly impact the size of the resulting NBs. However, non-pressurized size isolation does yield a significantly lower concentration of the isolated NBs. The process of centrifugation in a non-pressurized system does not significantly impact the ability to size isolate nanoscale agents, but it does result in NBs with a lower concentration which produce more variable nonlinear signal intensity compared to the pressurized control. The removal of the polydisperse bubble solution from the closed system saturated with C_3_F_8_ gas to the non-pressurized vessel exposed to air likely desaturates the system surrounding the NBs of PFC which increases the rate of PFC diffusion from the NB cores to the surrounding medium. The loss of C_3_F_8_ from the NB cores may reduce their stability thereby diminishing their ultrasound activity, as demonstrated by the reduction in signal intensity. Additionally, the increased handling of the material may negatively impact NB yield, contributing to the significant decrease in concentration observed in the non-pressuri ze d condition. These data suggest that the highest concentration, most echogenic NBs are produced when the NBs remain in the PFC saturated, pressurized vessel.

## Conclusions

The production of NBs with a uniform size distribution, high concentration and consistent echogenic properties requires careful production controls. This work identifies some critical processing steps in the production of NBs via mechanical amalgamation.

Characterization of the lipid dispersion precursor confirmed that sonication intensity alters the size distribution of precursor particles in the lipid dispersion. Confirmed through transmittance measurement and DLS sizing, the size distribution of precursor particles in the dispersion gets smaller and more uniform (multi-to monomodal, narrower distribution) as the sonication intensity increases. Despite this change, XPS analysis demonstrated that the elemental composition of all lipid dispersion precursors was not altered. Our data showed that the NBs produced from lipid dispersion which underwent varying sonication protocols had no significant difference in diameter or concentration. Despite this, the NBs produced from a lipid dispersion which underwent the highest intensity sonication had the most stable acoustic activity. This is likely due to the lipid dispersion precursor being homogenous and the most uniform in size, resulting in NBs with homogenous shell lipid distribution.^28^

In terms of gas to lipid ratio, we found that gas volume in excess of the lipid dispersion volume is required to produce high concentration, echogenic NBs. When the gas volume was equal to the lipid dispersion volume, the resulting NBs had significantly lower concentration and poor stability with and without the presence of US. The mixed gas (C_3_F_8_/atmosphericair) cores of these NBs are more susceptible to outward air diffusion, reducing their ability to produce ultrasound signal. Finally, we observed that size isolation in a pressurized vessel results in significantly higher NB concentration and increased stability under US compared to non-pressurized size isolation. The increased handling required in this protocol and exposure to air increases material loss and gas diffusion from NBs, decreasing the acoustic activity they are able to generate.

Our findings highlight the importance of lipid dispersion precursor homogeneity for NB stability under US, gas volume in excess of lipid dispersion volume for high concentration NB production, and efficient size isolation techniques for high yield of NBs. These results emphasize that the production of NBs requires careful production controls, however with the proper experimental design, it is possible to produce echogenic, submicron contrast agents repeatedly and reliably.

This work offers insight into how to design a mechanical amalgamation-based NB production method for the manufacturing of NBs for use as UCAs. These findings can be translated to improve MB (on-market UCAs) production as similar mechanical amalgamation techniques are used to produce these micron scale agents. Additionally, a thorough understanding of how manufacturing parameters affect agent size is beneficial for NBs in nonmedical applications (i.e. water treatment, agriculture).

Finally, this work seeks to shed light on the complexities of producing NBs by highlighting how the production of nanoscale, uniform diameter, high concentration, and critically, echogenic, NBs is susceptible to protocol deviations but with the proper controls in place, such protocols can be successfully executed.

## Author contributions

Theresa Kosmides: conceptualization, data curation, formal analysis, methodology, investigation, writing – original draft. Dana Wegierak: software, investigation, writing – review & editing. Aaqib H. Khan: investigation, writing – review & editing. Ilya Bederman: investigation. Tae Kyong John Kim: investigation. Agata Exner: supervision, resources, funding acquisition, writing – review & editing.

## Conflicts of interest

The authors declare the following competing financial interest(s): A.A.E. is the founder of Visano Theranostics.

## Data availability

The data supporting this article are not currently publicly available at this time but may be obtained from the authors upon reasonable request.

## Acknowledgements

This work was supported by the National Institutes of Health (R01EB025741). XPS was conducted at the Case School of Engineering’s Swagelok Center for Surface Analysis of Materials (CSE-SCSAM). Financial assistance for instrument time and scientific training was provided by the SCSAM Fellowship program which is supported by the SCSAM Endowment Fund. Figures were created by BioRender and GraphPad Prism. No artificial intelligence tools were used in the experimental design, data acquisition, analysis, or writing of this work.

## Notes

### Competing Interest Statement

The authors have declared no competing interest.

## Notes and references

1. Lasek L, Krzywanski J, Skrobek D, Zylka A, Nowak W. Review of Micro- and Nanobubble Technologies: Advancements in Theory and Applications and Perspectives on Adsorption Cooling and Desalination Systems. Energies. 2023;16(24):8078. doi:10.3390/en16248078

2. Lohse D, Zhang X. Surface nanobubbles and nanodroplets. Rev Mod Phys. 2015 Aug 31;87(3):981–1035. doi:10.1103/RevModPhys.87.981

3. Unger EC, Porter T, Culp W, Labell R, Matsunaga T, Zutshi R. Therapeutic applications of lipid-coated microbubbles. Adv Lipid-Based Drug Solubilization Target. 2004 May 7;56(9):1291–314. doi:10.1016/j.addr.2003.12.006

4. Fine bubble technology — General principles for usage and measurement of fine bubbles — Part 1: Terminology [Internet]. ISO 20480-1:2017. Available from: https://www.iso.org/obp/ui/#iso:std:iso:20480:-1:ed-1:v1:en

5. Jia M, Farid MU, Kharraz JA, Kumar NM, Chopra SS, Jang A, et al. Nanobubbles in water and wastewater treatment systems: Small bubbles making big difference. Water Res. 2023 Oct 15;245:120613. doi:10.1016/j.watres.2023.120613

6. Chen W, Bastida F, Liu Y, Zhou Y, He J, Song P, et al. Nanobubble oxygenated increases crop production via soil structure improvement: The perspective of microbially mediated effects. Agric Water Manag. 2023 May 31;282:108263. doi:10.1016/j.agwat.2023.108263

7. Hansen HHWB, Cha H, Ouyang L, Zhang J, Jin B, Stratton H, et al. Nanobubble technologies: Applications in therapy from molecular to cellular level. Biotechnol Adv. 2023 Mar 1;63:108091. doi:10.1016/j.biotechadv.2022.108091

8. Ramirez DG, Abenojar E, Hernandez C, Lorberbaum DS, Papazian LA, Passman S, et al. Contrast-enhanced ultrasound with sub-micron sized contrast agents detects insulitis in mouse models of type1 diabetes. Nat Commun. 2020 May 7;11(1):2238. doi:10.1038/s41467-020-15957-8

9. Yano Y, Hamano N, Haruta K, Kobayashi T, Sato M, Kikkawa Y, et al. Development of an Antibody Delivery Method for Cancer Treatment by Combining Ultrasound with Therapeutic Antibody-Modified Nanobubbles Using Fc-Binding Polypeptide. Pharmaceutics. 2023;15(1). doi:10.3390/pharmaceutics15010130

10. Nittayacharn P, Abenojar E, Cooley MB, Berg FM, Counil C, Sojahrood AJ, et al. Efficient ultrasound-mediated drug delivery to orthotopic liver tumors – Direct comparison of doxorubicin-loaded nanobubbles and microbubbles. J Controlled Release. 2024 Mar 1;367:135–47. doi:10.1016/j.jconrel.2024.01.028

11. Frinking P, Segers T, Luan Y, Tranquart F. Three Decades of Ultrasound Contrast Agents: A Review of the Past, Present and Future Improvements. Ultrasound Med Biol. 2020 Apr 1;46(4):892–908. doi:10.1016/j.ultrasmedbio.2019.12.008

12. Paefgen V, Doleschel D, Kiessling F. Evolution of contrast agents for ultrasound imaging and ultrasound-mediated drug delivery. Front Pharmacol. 2015;Volume 6–2015. doi:10.3389/fphar.2015.00197

13. McArthur C, Baxter GM. Current and potential renal applications of contrast-enhanced ultrasound. Clin Radiol. 2012 Sep 1;67(9):909–22. doi:10.1016/j.crad.2012.01.017

14. Gao Y, Hernandez C, Yuan HX, Lilly J, Kota P, Zhou H, et al. Ultrasound molecular imaging of ovarian cancer with CA-125 targeted nanobubble contrast agents. Nanomedicine Nanotechnol Biol Med. 2017 Oct 1;13(7):2159–68. doi:10.1016/j.nano.2017.06.001

15. Exner AA, Kolios MC. Bursting microbubbles: How nanobubble contrast agents can enable the future of medical ultrasound molecular imaging and image-guided therapy. Curr Opin Colloid Interface Sci. 2021;54:101463. doi:10.1016/j.cocis.2021.101463 PubMed PMID: 34393610 PMCID: PMC8356903.

16. Berg FM, Abenojar EC, Nittayacharn P, Hoggard NK, Tavri S, Wang J, et al. Nanobubbles for Precision Oncology: Preclinical Evaluation of Molecular-Targeted Ultrasound Contrast Agents in a Rabbit Model. bioRxiv. 2025. doi:10.1101/2025.08.07.668901

17. Yu H, Zheng S, Wang C, Xing J, Li L. Novel anti-VEGFR2 antibody-conjugated nanobubbles for targeted ultrasound molecular imaging in a rabbit VX2 hepatic tumor model. J Mater Chem B. 2023;11(45):10956–66. doi:10.1039/D3TB01718D

18. Nittayacharn P, de Leon A, Abenojar EC, Exner AA. The Effect of Lipid Solubilization on the Performance of Doxorubicin-Loaded Nanobubbles. In: 2018 IEEE International Ultrasonics Symposium (IUS). 2018. p. 1–4. doi:10.1109/ULTSYM.2018.8579716

19. Prabhakar A, Banerjee R. Nanobubble Liposome Complexes for Diagnostic Imaging and Ultrasound-Triggered Drug Delivery in Cancers: A Theranostic Approach. ACS Omega. 2019 Sep 24;4(13):15567–80. doi:10.1021/acsomega.9b01924

20. Yasuda K, Matsushima H, Asakura Y. Generation and reduction of bulk nanobubbles by ultrasonic irradiation. Chem Eng Sci. 2019 Feb 23;195:455–61. doi:10.1016/j.ces.2018.09.044

21. Peyman SA, McLaughlan JR, Abou-Saleh RH, Marston G, Johnson BRG, Freear S, et al. On-chip preparation of nanoscale contrast agents towards high-resolution ultrasound imaging. Lab Chip. 2016;16(4):679–87. doi:10.1039/C5LC01394A

22. Counil C, Abenojar E, Perera R, Exner AA. Extrusion: A New Method for Rapid Formulation of High-Yield, Monodisperse Nanobubbles. Small. 2022;18(24):2200810. doi:10.1002/smll.202200810

23. de Leon A, Perera R, Hernandez C, Cooley M, Jung O, Jeganathan S, et al. Contrast enhanced ultrasound imaging by nature-inspired ultrastable echogenic nanobubbles. Nanoscale. 2019;11(33):15647–58. doi:10.1039/C9NR04828F PubMed PMID: 31408083 PMCID: PMC6716144.

24. Frinking P, Segers T, Luan Y, Tranquart F. Three Decades of Ultrasound Contrast Agents: A Review of the Past, Present and Future Improvements. Ultrasound Med Biol. 2020;46(4):892– 908. doi:10.1016/j.ultrasmedbio.2019.12.008

25. Kaya M, Gregory V TS, Dayton PA. Changes in Lipid-Encapsulated Microbubble Population During Continuous Infusion and Methods to Maintain Consistency. Ultrasound Med Biol. 2009 Oct 1;35(10):1748–55. doi:10.1016/j.ultrasmedbio.2009.04.023

26. Abdalkader R, Kawakami S, Unga J, Higuchi Y, Suzuki R, Maruyama K, et al. The development of mechanically formed stable nanobubbles intended for sonoporation-mediated gene transfection. Drug Deliv. 2017 Nov;24(1):320–7. doi:10.1080/10717544.2016.1250139 PubMed PMID: 28165819; PubMed Central PMCID: PMC8241156.

27. Sirsi S, Borden M. Microbubble Compositions, Properties and Biomedical Applications. Bubble Sci Eng Technol. 2009 Nov;1(1–2):3–17. doi:10.1179/175889709X446507 PubMed PMID: 20574549; PubMed Central PMCID: PMC2889676.

28. Langeveld SAG, Schwieger C, Beekers I, Blaffert J, van Rooij T, Blume A, et al. Ligand Distribution and Lipid Phase Behavior in Phospholipid-Coated Microbubbles and Monolayers. Langmuir. 2020 Mar 31;36(12):3221–33. doi:10.1021/acs.langmuir.9b03912

