## Supplemental Figure 1 for "Critical Process Steps for Mechanical Agitation Driven Coated Nanobubble Self Assembly"

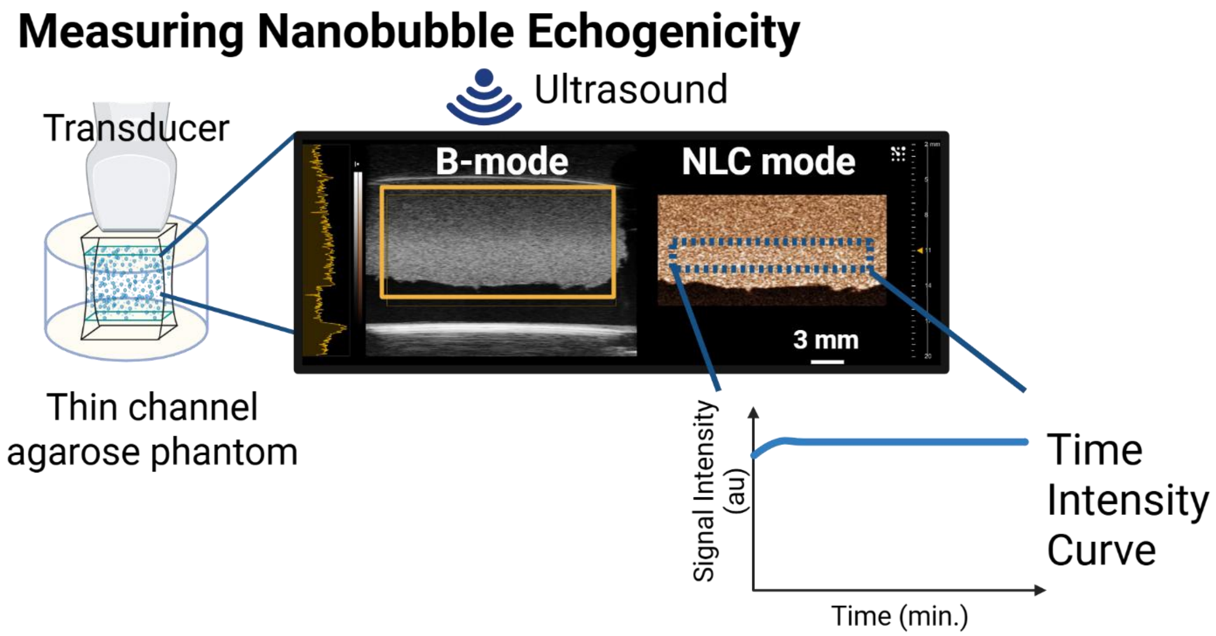


Fig S1. Overview of quantification of nanobubble echogenicity. Nanobubbles are diluted with PBS and placed in an agarose phantom with a channel of equivalent dimensions to the transducer (22 x 1 x 10 mm). The ultrasound transducer is directly coupled to the nanobubble suspension. Ultrasound data is collected in dual B mode and nonlinear contrast (NLC) mode. A region of interest (ROI) is drawn at the focal depth and the average NLC signal as a function of time is calculated for the ROI.
